# Early Detection of Apathetic Phenotypes in Huntington’s Disease Knock-in Mice Using Open Source Tools

**DOI:** 10.1101/208520

**Authors:** Shawn Minnig, Robert M. Bragg, Hardeep S. Tiwana, Wes T. Solem, William S. Hovander, Eva-Mari S. Vik, Madeline Hamilton, Samuel R. W. Legg, Dominic D. Shuttleworth, Sydney R. Coffey, Jeffrey P. Cantle, Jeffrey B. Carroll

## Abstract

Apathy is one of the most prevalent and progressive psychiatric symptom in Huntington’s disease (HD) patients. However, preclinical work in HD mouse models tend to focus on molecular and motor, rather than affective, phenotypes. Measuring behavior in mice often produces noisy data and requires large cohorts to detect phenotypic rescue with appropriate power. The operant equipment necessary for measuring affective phenotypes is typically expensive, proprietary to commercial entities, and bulky which can render adequately sized mouse cohorts as cost-prohibitive. Thus, we describe here a home-built open-source alternative to commercial hardware that is reliable, scalable, and reproducible. Using off-the-shelf hardware, we adapted and built several of the rodent operant buckets (ROBucket) designed to test *Htt^Q111/+^* mice for attention deficits in fixed ratio (FR) and progressive ratio (PR) tasks. We find that, despite normal performance in reward attainment in the FR task, *Htt^Q111/+^* mice exhibit reduced PR performance at 9-11 months of age, suggesting motivational deficits. We replicated this in two independent cohorts, which demonstrates the reliability and utility of both the apathetic phenotype, and these ROBuckets, for preclinical HD studies.

## Introduction

Huntington’s disease (HD) is a fatal, progressive neurological disorder caused by a coding cytosine-adenine-guanine (CAG) expansion in the huntingtin (*HTT*) gene, where repeat lengths above 40 result in full penetrance of the disease ^1^. Patients with HD display a clinical triad of cognitive, psychiatric, and motor symptoms ^2^. Although chorea - a hyperkinetic movement disorder consisting of brief, irregular movements used by clinicians as a prerequisite for formal clinical diagnosis ^3,4^ - is the most recognized HD related deficit ^5^, cognitive and psychiatric symptoms often appear during the prodromal phase of the disease, as many as 10 years prior to the onset of motor dysfunction ^6,7^. Cognitive symptoms in HD include various impairments in learning and memory, deficits in executive functioning ^8^, and difficulty recognizing emotional states ^9,10^, while the most commonly identified psychiatric manifestations include apathy, anxiety, depression, irritability, perseveration, and obsessive behaviors ^6,7,11^.

The cognitive and behavioral aspects of HD contribute to significant declines in functional capacity (i.e. the activities of everyday life) ^12^, and are often described as being the most burdensome symptoms to both HD patients and their families ^13^. In the absence of effective disease modifying treatments, current therapeutic options for HD are focused on managing these symptoms ^14^. As such, a great need exists to further study the cognitive and psychiatric components of HD, not only because of the great distress they cause to HD families, but also because understanding the progression of these symptoms will provide researchers with the opportunity to assess the efficacy of potential disease modifying therapies at the earliest time point possible. Unfortunately, given the importance of these phenotypes in HD patients’ quality of life, most HD preclinical mouse studies do not typically include analysis of impaired cognition and altered affect ^15^. We believe one reason for this is the expense and complexity of the equipment required for traditional cognitive and affective assays, limiting the number of animals feasible to study in preclinical HD studies. This practical limitation of study size limits the power of HD preclinical studies which, as is common in neuroscience research ^16^, is generally poor.

Amongst HD’s psychiatric manifestations, apathy has an extremely high point prevalence, coupled with a uniquely consistent relationship between severity and HD progression ^7,17^. A recent large study of presymptomatic HD mutation carriers found striking increases in the incidence of apathy, even in subjects calculated to be more than 10 years from clinical onset ^18^. Apathy, as a psychiatric symptom distinct from depression, has been operationalized to contain aspects of diminished motivation, reduced goal-directed behavior, lack of interest in new experiences, and diminished emotional responsivity ^19^. In HD patients, apathy is correlated with functional capacity and cognition, but not depression, suggesting apathy is a distinctive component of the affective landscape of HD ^20^.

Motivated by the importance of apathy to the lived experience of HD mutation carriers, we are interested in bringing analysis of apathy into HD preclinical studies. Traditional rodent experiments to test motivated behavior include the progressive-ratio (PR) operant task ^21^, in which subjects are required to perform increasingly large numbers of nose-pokes or lever-presses to receive a reward. Operant chambers used to assay PR responses in rodents can cost thousands of dollars each, limiting the number of animals (and thereby statistical power) of preclinical studies of apathy. To address this problem, we have modified a recently described open-source operant chamber - the “ROBucket” ^22^ - based on the Arduino computing platform, with a total built cost of approximately $150/chamber. To validate the modified apparatus, we studied motivation in 9-11 month old B6.*Htt*^Q*111*/+^ mice, a knock-in mouse model of the HD mutation ^23^. Using commercially available tools we, and others, have previously observed motivational phenotypes in this model that precede motor or cognitive changes ^24–27^. Using the open-source ROBucket, we confirm specific deficits in progressive, but not fixed, ratio tasks in B6.*Htt*^Q*111*/+^ mice at this time point, consistent with relatively intact learning but impaired motivation.

## Results

### Modification of existing operant chamber design

We first precisely recreated a recently reported open source operant chamber (“ROBucket”) ^22^. On conducting pilot experiments, we found mice tended to interact with the housing and reward tubing fed through the bucket in the original design. To overcome this distraction, we redesigned the 3D-housing apparatus to be outside the mouse chamber (Fig. 1) - updated plans are available online at https://zenodo.org/record/1011360 ^28^. Studies performed with these modifications revealed that mice quickly learned to interact with the active well to receive 10 μl of 20% sucrose, and that the modified ROBuckets (mROBucket) accurately counted nose pokes, compared to direct observations.

**Fig 1.**
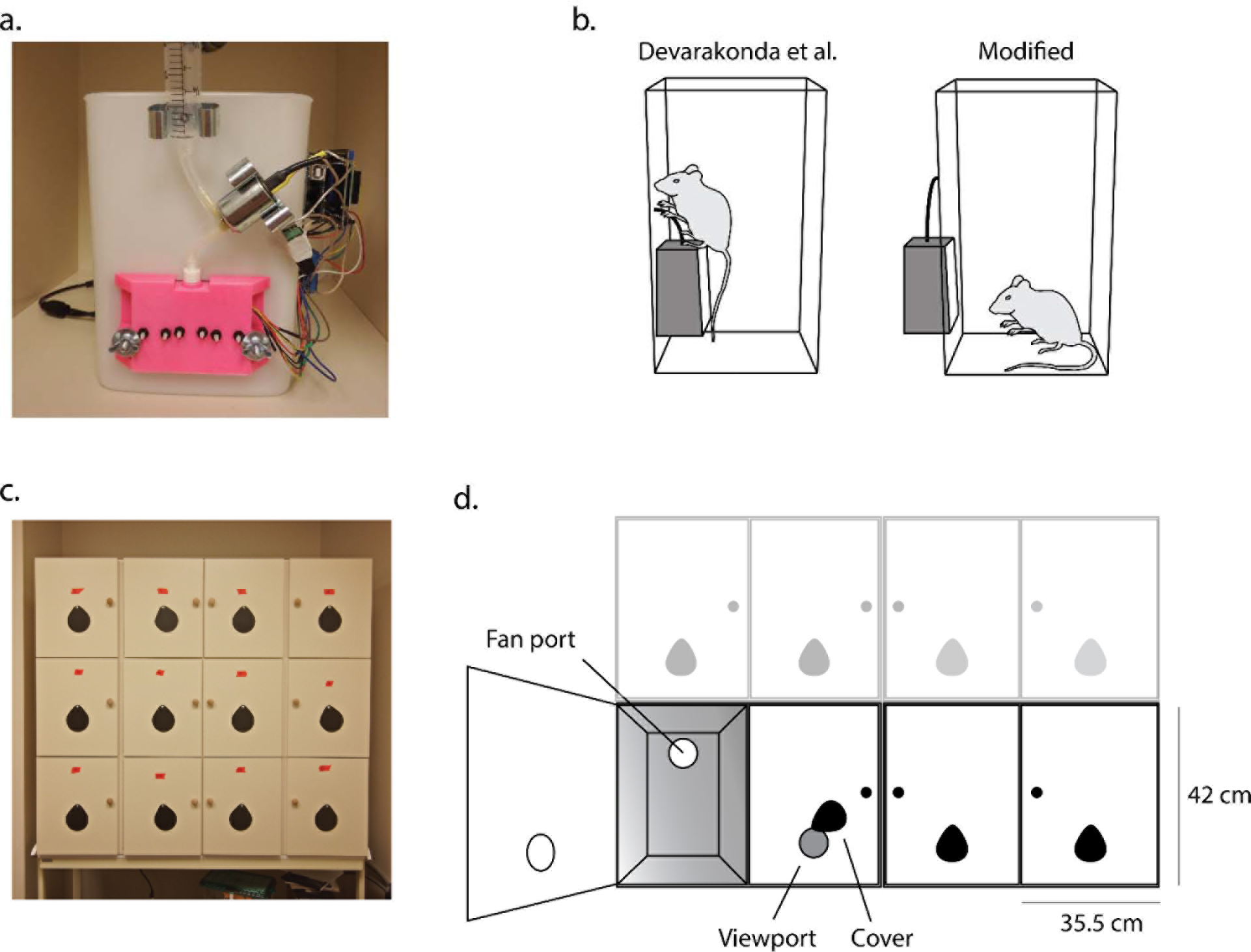
Modified ROBucket design and housing. a-b. mROBucket with nose-poke housing mounted on the exterior of the bucket reduced time spent exploring the housing and sucrose tubing. **c-d**. Construction of a 4×3 grid of isolation housing was used to keep each mROBucket independent and facilitate concurrent testing of adequately sized cohorts. Each housing chamber included a small fan for white noise and a viewport to monitor the mROBucket display throughout the experiment.

### Normal fixed ratio performance in *Htt*^Q*111*/+^ mice

9-month-old male B6.*Htt*^+/+^ (*n* = 7) and B6.*Htt^Q111/+^* (*n* = 12) mice (hereafter *Htt*^+/+^ and *Htt^Q111/+^*) were single-housed and food restricted over 2 weeks (target weight loss of 2% / day, final body weight ~85% free-feeding weight) before operant testing. We observed no impact of genotype on baseline body weight or the rate at which *Htt*^+/+^ and *Htt^Q111/+^* mice lost weight during food restriction (Supplemental Fig 1). After body weight stabilized, mice were placed into mROBucket chambers for 1 hour sessions each day on a fixed ratio 1 (FR1) reinforcement schedule - i.e. one nose poke in the active well resulted in sucrose delivery in the reward well. There was a 1 second timeout after each active well response. Two criteria were required for progression to the next phase: a 3:1 preference for the active well versus the inactive well and 20 or more reinforcements for 3 consecutive days. All mice, with the exception of a single *Htt*^+/+^ mouse (Fig. 2, grey panel), quickly learned the task (Fig. 2, average time to FR1 criteria 7.5 ± 2.4 days).

**Fig 2.**
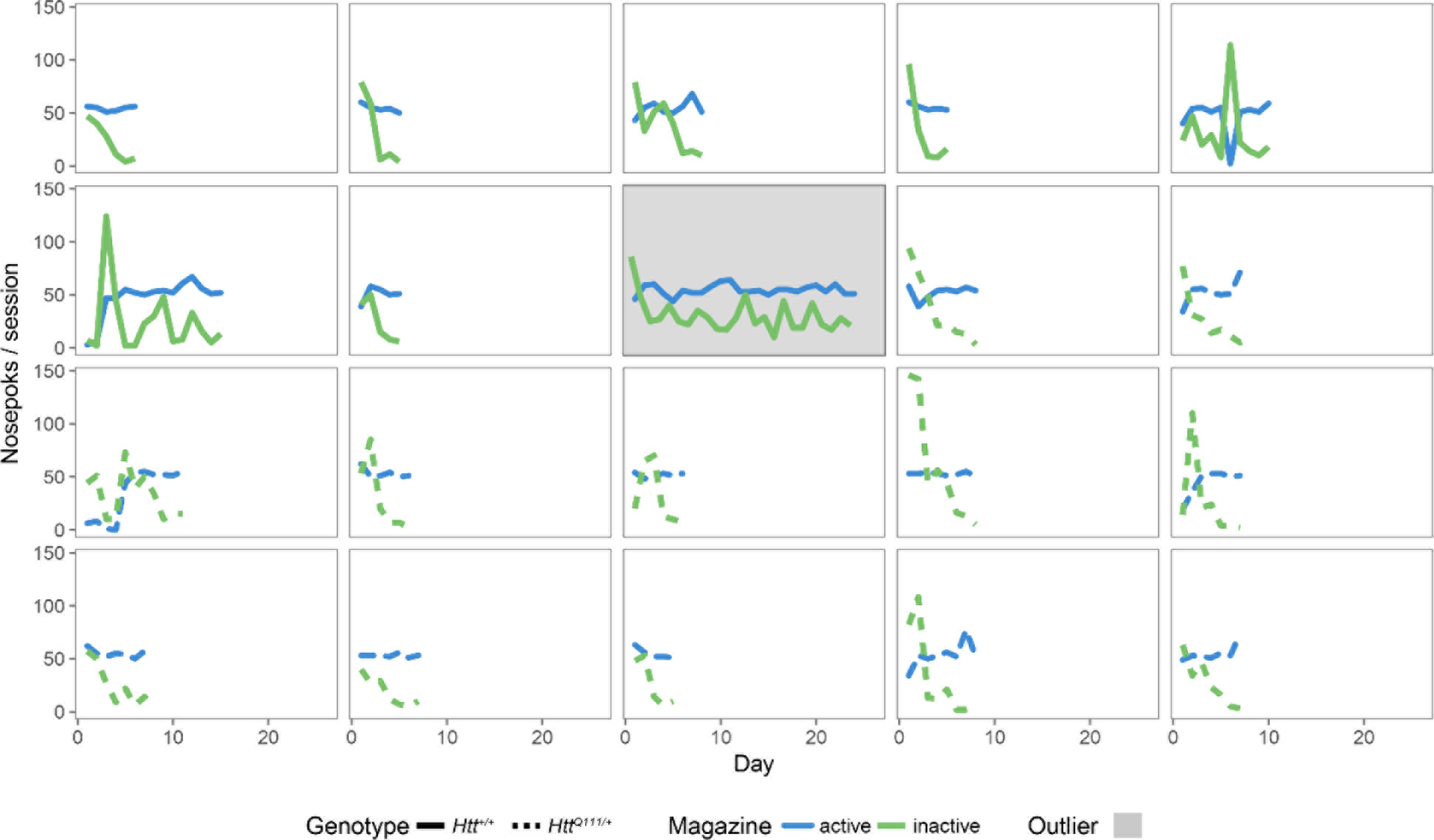
Normal per-mouse acquisition of FR1 task in 10-month-old *Htt^Q111/+^* mice. Shown for each mouse is the number of nose pokes per 1 hour session in the active (blue) and inactive (green) wells. *Htt*^+/+^ mice (*n* = 8) are graphed with solid lines, *Htt^Q111/+^* with dashed lines (*n* = 12). One mouse (grey highlight) was excluded for failing to pass the pre-defined FR1 acquisition criteria.

We observed no effect of genotype on days to meet criteria, the active/inactive nose poke ratio, or total nose pokes per session on the FR1 task (Fig. 3). This suggests 9-month-old male *Htt^Q111/+^* mice are able to normally acquire this simple discrimination task.

**Fig 3.**
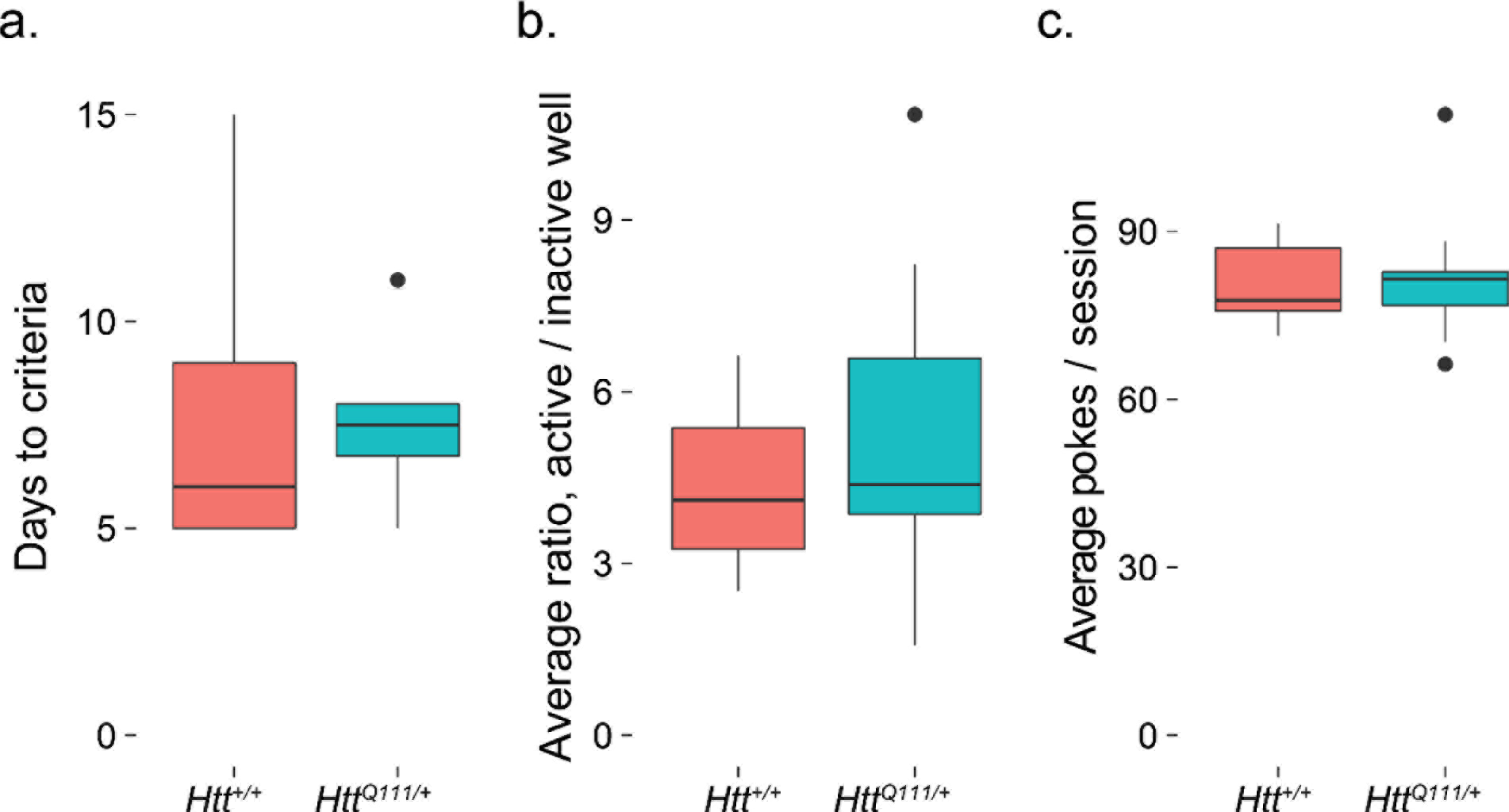
Normal performance of FR1 task in 10-month-old male *Htt^Q111/+^* mice. No genotype effect is observed in the days to criteria (**a.** *t*_(7.1)_ = 0.2, *p* = 0.8), active/inactive well ratio (**b.** t_(16.9)_ = -0.9, p = 0.4) or average total nose pokes per session (**c.** *t*_(16.5)_ = -0.1, *p* = 0.9). Horizontal lines in the boxes indicate 25%, 50% and 75% percentiles, while vertical lines indicate 1.5 times the interquartile range; outliers beyond these values are graphed as points.

After meeting both criteria for FR1, we next trained the mice on a fixed ratio 5 (FR5) task for 3 days to familiarize them with tasks requiring multiple nose pokes to achieve reward. There was a 1 second timeout after each active well response. Consistent with the FR1 task, 10-month-old male *Htt^Q111/+^* mice are able to normally acquire this simple discrimination task, with no differences observed between the ratio of active/inactive well responses or the total number of nose pokes per session (Fig. 4). This suggests fatigue and motor dysfunction do not prevent *Htt^Q111/+^* mice from making large numbers of accurate nose pokes in an hour-long session(average of 273.1 ± 11.7 nose pokes/session).

**Fig 4.**
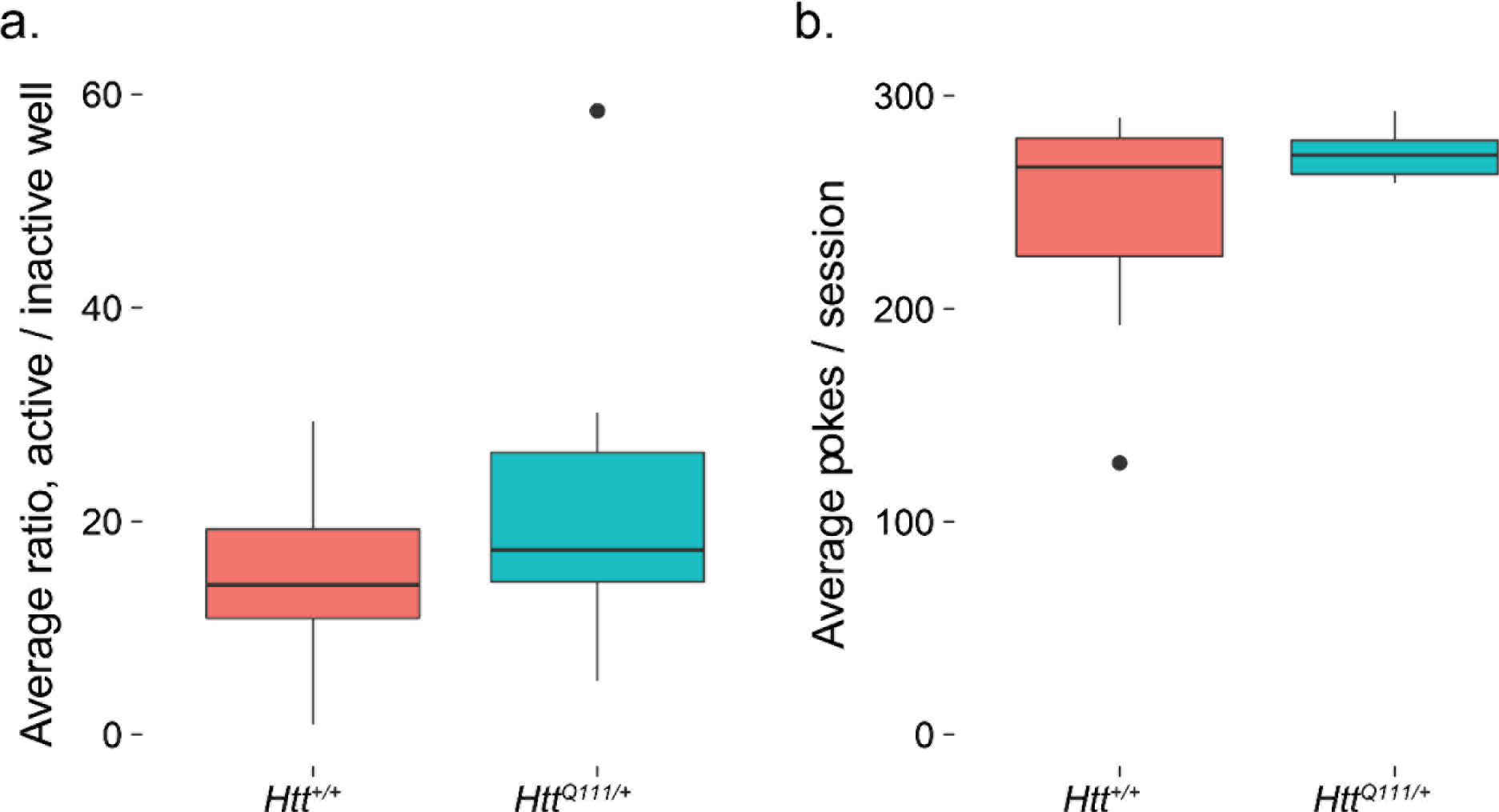
Normal performance of FR5 task in 10-month-old *Htt^Q111/+^* mice. No genotype effect is observed in the active/inactive well ratio (**a.** *t*_(16.6)_ = -1.3, *p* = 0.2) or average total nose pokes per session (**b.** *t*_(6‥3)_ = -1.4, *p* = 0.2); *n* = 7 *Htt*^+/+^, 12 *Htt^Q111/+^*. Horizontal lines in the boxes indicate 25%, 50% and 75% percentiles, while vertical lines indicate 1.5 times the interquartile range; outliers beyond these values are graphed as points.

### Reduced progressive ratio performance in *Htt^Q111/+^* mice

Regardless of performance, mice were advanced to a progressive ratio (PR) task after 3 days of FR5 training. This task requires the animals to respond with an exponentially increasing number of sequential nose pokes in the active well to receive the reward (1, 2, 4, 6, 9, 12, etc., according to the formula R = ‖5e^(N*0.2)^‖ – 5)^29^. The final number of reinforcements achieved is referred to as the “breakpoint” ^21^. We established the criterion for the PR task based on breakpoint stabilization, or less than 10% variation in the breakpoint in 3 consecutive trials. Both *Htt*^+/+^ and *Htt^Q111/+^* mice learned the PR task, with breakpoints stabilizing between 4-17 days.

We observed no genotype effect on the days to criterion (Fig. 5a) or average active/inactive ratio (Fig. 5b) during the PR task, however *Htt^Q111/+^* mice perform significantly fewer total nose pokes per session (44% reduction, Fig. 5c) and consequently receive fewer rewards per session (17% reduction). This results in a 37% reduction in the final stabilized breakpoint of *Htt^Q111/+^* mice compared to *Htt*^+/+^ mice (Fig. 5d). Post-hoc analysis suggests our experiment had 94.6% power to detect a breakpoint difference between genotypes (*n* = 7 *Htt*^+/+^ and 12 *Htt^Q111/+^*; effect size, *d*, = 1.8, type 1 error probability = 0.05).

**Fig 5.**
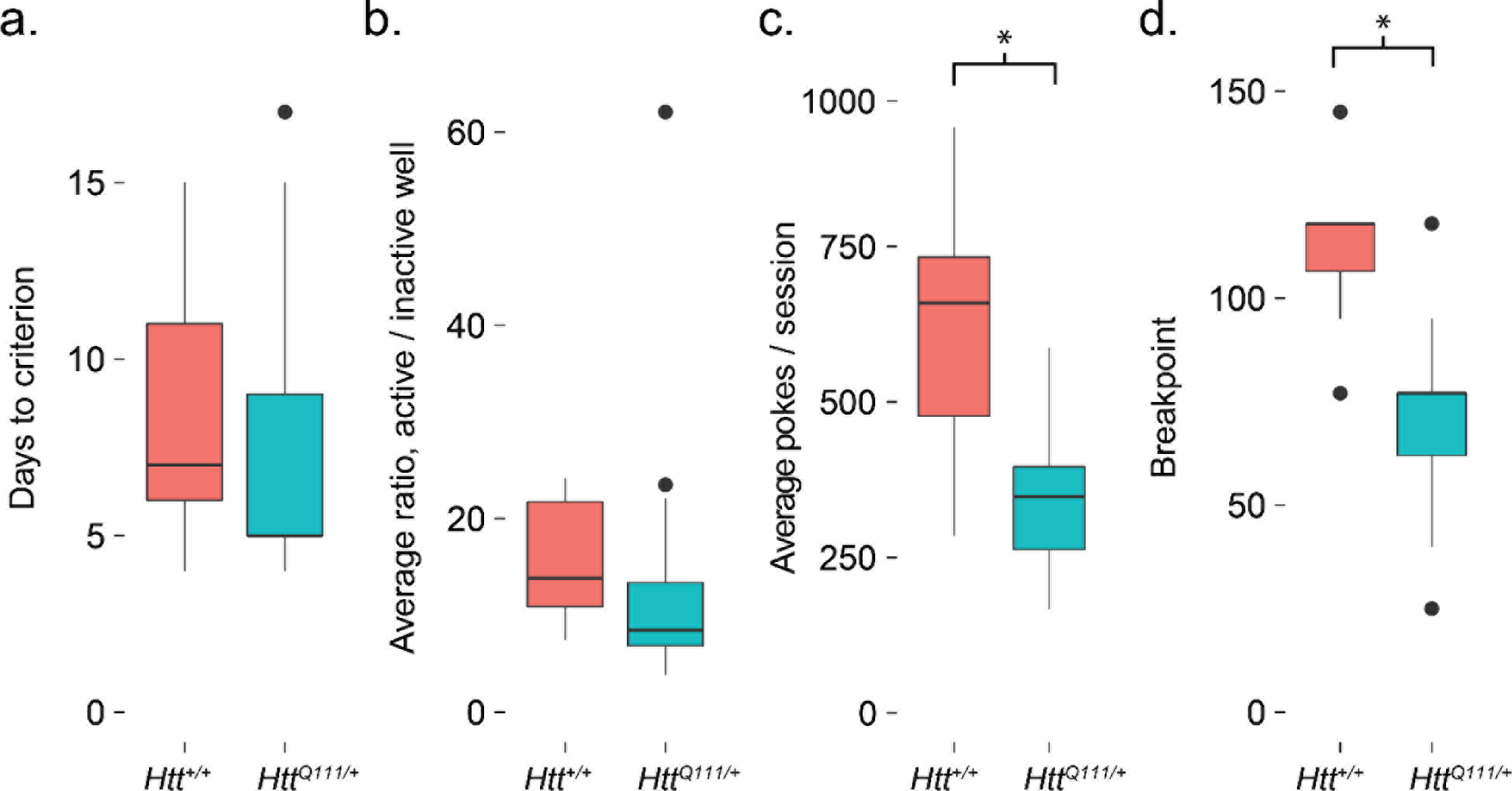
Progressive ratio deficits in 10-month-old Htt^+/+^ and *Htt^Q111/+^* mice. No genotype effect is observed in (a.) the final days to criterion (*t*_(14.3)_ = 0.5, *p* = 0.6), or (b.) active/inactive well ratio ( *t*_(15.8)_ = 0.3, *p* = 0.8) during the PR task. However, *Htt^Q111/+^* mice do have (c.) reduced total nose pokes (*t*_(7.8)_ = 3.0, *p* = 0.02), reduced rewards/session (*t*_(12.1)_ = 3.0, *p* = 0.01) and a consequently (d.) reduced final breakpoint (*t*_(13.9)_ = 4.0, *p* = 0.002) during the PR task. * indicates *p* < 0.05. Horizontal lines in the boxes indicate 25%, 50% and 75% percentiles, while vertical lines indicate 1.5 times the interquartile range; outliers beyond these values are graphed as points.

### Replication in an independent cohort

We are interested in the improvement of preclinical trials in HD, particularly their reproducibility ^27,30-32^,. To determine whether the mROBucket apparatus and progressive ratio task are reproducible assays of motivated behavior, we repeated the experiment in a new cohort of similarly aged (10-11 months) male *Htt*^+/+^ (*n* = 12) and *Htt^Q111/+^* mice (n = 12). Consistent with the findings of our first cohort, we observe no effect of genotype on the total number of nose pokes in the FR1 task (Fig. 6a). Similarly, in the FR5 task, *Htt^Q111/+^* mice perform equivalently to *Htt*^+/+^ mice for total nose pokes (Fig. 6b). In the PR task, *Htt^Q111/+^* displayed reduced total pokes (*t*_(17.1)_ = 3.9, *p* = < 0.001), earned fewer rewards (*t*_(21.5)_ = 3.9, *p* < 0.001) and displayed a 46% reduction in final breakpoint (Fig. 6c). Post-hoc analysis of the replication cohort (*n* = 12 *Htt*^+/+^ and 12 *Htt^Q111/+^*, effect size, *d*, = 1.6, type 1 error probability = 0.05) indicates we achieved 96.3% power.

**Fig 6.**
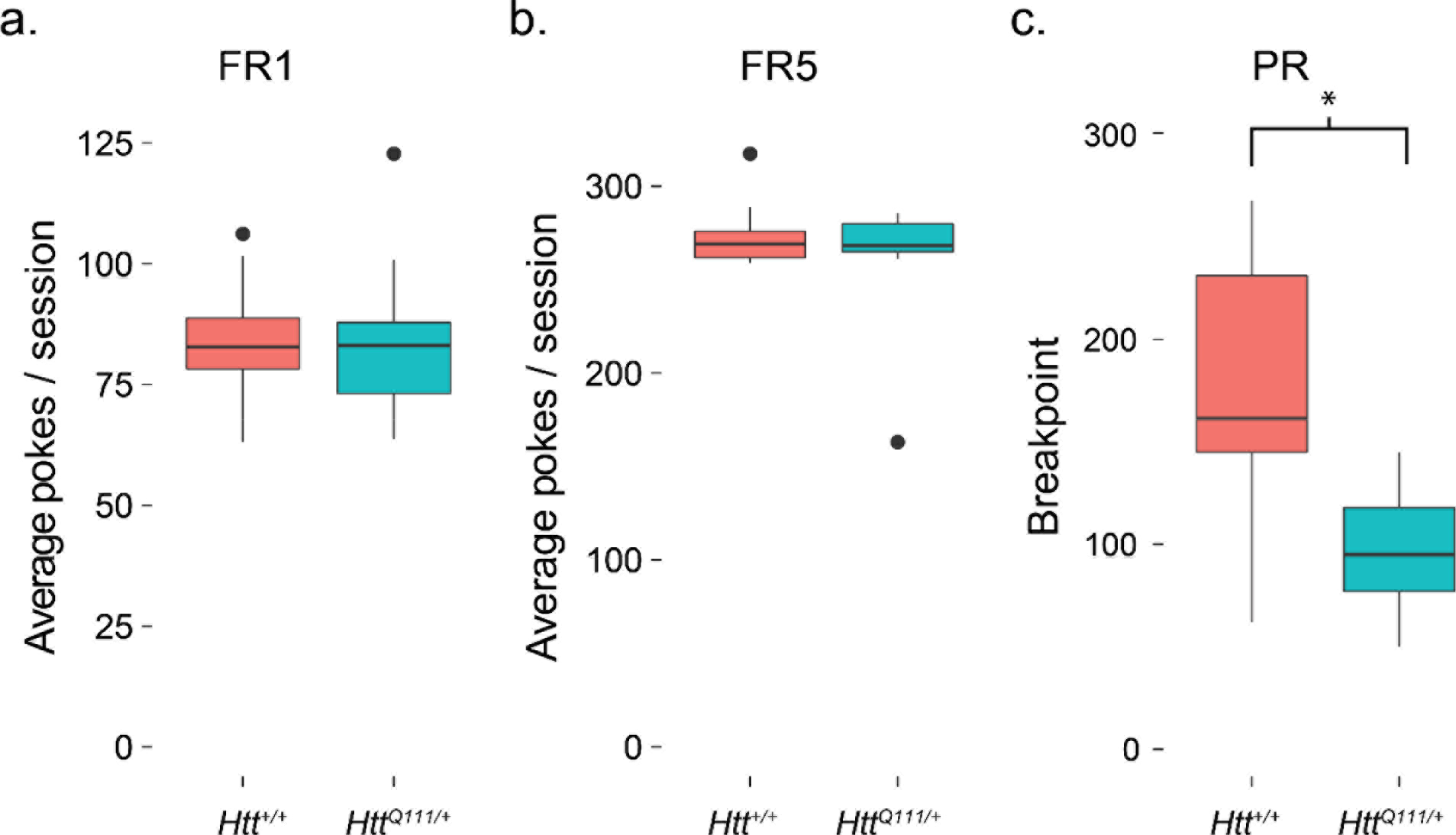
Independent replication of progressive ratio deficits in 10-month-old *Htt*^+/+^ and *Htt^Q111/+^* mice. **a.** We observed no difference in the total nose pokes during the FR1 task (*t*_(19.8)_ = 0.12, *p* = 0.9). **b.** Similarly, in the FR5 task, *Htt^Q111/+^* mice perform equivalently to *Htt*^+/+^ mice (*t*_(16.3)_ = 0.9, *p* = 0.4). **c.** The replication cohort had a significantly reduced final breakpoint (*t*_(16.2)_ = 3.9, *p* = 0.001) in the PR task. * indicates *p* < 0.05. Horizontal lines in the boxes indicate 25%, 50% and 75% percentiles, while vertical lines indicate 1.5 times the interquartile range; outliers beyond these values are graphed as points.

## Discussion

Apathy is a core feature of affective dysfunction in HD ^17^, but analysis of apathy and other affective disturbances is rarely included in preclinical studies of HD ^33^. Here, we demonstrate that an inexpensive open source apparatus can robustly and reproducibly detect motivational deficits in 9-11 month old *Htt^Q111/+^* mice. These motivational deficits precede overt motor, cognitive, or neurodegenerative changes in this model ^27^, suggesting they may occur amongst the earliest changes associated with mutant huntingtin expression *in vivo*.

The basal ganglia, and particularly the striatum, are the most strikingly impacted brain regions in Huntington’s disease, showing robust volume declines many years before clinical disease onset ^7,34^. While no HD mouse models experience similar pronounced neurodegeneration, a number of imaging ^30,35,36^ and molecular analyses ^37^ confirm that the caudoputamen is the most strikingly impacted brain region in mice expressing mutant *HTT*. In humans, focal ischemic basal ganglia damage is associated with a range of motivational deficits ^38^, including notable deficits in incentive motivation - the process of activating specific behavioral responses based on predicted reward ^39^. In stroke patients with basal ganglia lesions, deficits in incentive motivation occur in the absence of deficits in hedonic responses, consistent with reduced progressive ratio performance in *Htt^Q111/+^* mice (here, and ^24,27^) at a time when sucrose preference tasks suggest no alterations in hedonic drive in these mice ^27^. Changes in instrumental motivation may therefore serve as a translatable readout of basal ganglia dysfunction in mice.

Within the basal ganglia of HD mutation carriers, distinctive patterns of atrophy are observed, such that medial paraventricular caudate is amongst the earliest regions to experience neurodegeneration and gliosis ^40^. Shape analysis of subcortical structures in HD mutation carriers in the TRACK-HD study confirm that the medial body of the caudate is the structure with the earliest and greatest extent of atrophy in HD mutation carriers, even before symptom onset ^41^. Non-human primate studies suggest these periventricular regions of the caudate receive dense projections from regions within the orbital and medial prefrontal cortex (OMPFC) that comprise the “medial” OMPFC network ^42^. These PFC-basal ganglia loops have been proposed to play key roles in regulating mood and motivated behavior ^43^. In healthy humans, the personality trait of persistence predicts activity levels in the medial network of the OMPFC and the medial-ventral striatum ^44^. If the earliest degenerative changes in the HD brain occur in this OMPFC-medial caudate network, altered motivation may be one of the earliest readouts of basal ganglia dysfunction.

Statistical power in neuroscience studies, including behavioral studies, is generally very low ^16^, which has led to widespread calls for improvements in conducting and reporting preclinical studies ^45^. In HD specifically, positive mouse studies of creatine ^46^, minocycline ^47^ and coenzyme-Q10 ^48^ provided partial justification for large human studies of each compound, all of which were unsuccessful when tested in patients ^49-51^. The only behavioral endpoint used in the these specific trials was accelerating rotarod - by far the most commonly reported behavioral measure across HD mouse model studies ^33^. We propose that the use of endpoints more relevant to human HD patients, such as apathy, may improve the likelihood an intervention will translate from HD mouse models to HD patients.

One factor limiting the more widespread use of operant based cognitive and affective tasks in HD mouse models is the complexity and cost of the apparatus required. Traditional operant chambers from commercial suppliers cost thousands of dollars, placing a practical limit on the number of mice that can be screened in parallel in a single lab and thereby limiting the power of these studies. Recent technological developments - including the rapid development of inexpensive additive manufacturing (i.e. “3D printing”) and low-cost open source computing platforms (e.g. Raspberry Pi and Arduino) - enable open source alternatives to commercial products. Further, they allow for rapid design and software modifications to assay behavior through multiple paradigms. These tools have been applied to video tracking of behavior ^52^, integrated microscopy, temperature control, and optogenetics in small animal experiments ^53^, as well as the rodent operant chambers used here ^22^. These technologies render practical the fabrication of a large number of chambers (12 were used in the current study), so dozens of mice can be tested in parallel in a single lab, with a limited number of handlers.

These experiments confirm previous findings that motivational deficits occur before pronounced neurodegeneration in the *Htt^Q111/+^* model of HD. They also show that these deficits are assayable using inexpensive hardware and open source software tools, which should enable their more widespread utilization in preclinical studies in HD.

## Methods

### Mice

B6.*Htt^Q111^* mice, which have been previously described ^54^, were originally obtained from JAX (Research Resource Identifier: IMSR\_JAX:003456) and bred and maintained at the Western Washington University vivarium. Mice were group housed until 9-10 months of age and given access to food and water ad libitum, until two weeks before testing was to begin. For genotyping, presence or absence of the mutant allele was determined by polymerase chain reaction of gDNA using the primers CAG1 (5’-ATGAAGGCCTTCGAGTCCCTCAAGTCCTTC-3’)^55^ and HU3 (5’-GGCGGCTGAGGAAGCTGAGGA-3’) ^56^. All experiments were conducted in accordance to the NIH Guide for the Care and Use of Laboratory Animals and approved by the Western Washington University animal care and use committee (protocol 16-007).

### Apparatus

2-choice operant chambers were constructed according to the design from Devarakonda et al. (2016) with modifications. Briefly, these are Arduino-controlled operant boxes which deliver a liquid sucrose reward in response to nose-pokes with various reinforcement schedules. The apparatus here was modified to place the nose-poke photo-beam housing on the outer, rather than inner, wall of the operant chamber (see Fig. 1b). This eliminated time spent exploring the nose-poke housing and associated tubing, and resulted in cleaner acquisition of the FR/PR tasks. Isolation housing was designed and constructed as a grid of 3×4 chambers (35 × 42 × 42 cm each) with view ports in the front to visualize the response readings and vent fans in the back to produce white noise (Fig. 1d).

### Behavioral Testing

Mice were single house and fasted over two weeks to reduce body weight to 85% of free feeding weight prior to testing and maintained at this weight for the duration of testing. For FR1 testing, mice learned to nose-poke on a fixed-ratio reinforcement schedule where a single nose-poke in the active well elicits delivery of a sucrose reward (10 uL, 20% sucrose), with a 1-second timeout after each active well press. Trial duration was 60 minutes or until the subject received 50 reinforcements, at which point mice were promptly removed. Acquisition criteria for the FR1 schedule were met when mice exhibited discrimination criteria of ≥3:1 for the active:inactive well and received ≥ 20 reinforcements for 3 consecutive days. Mice not meeting the FR1 acquisition criteria by 17 days were excluded from further testing (this occurred in 1 mouse from each cohort, or approximately 5% of mice). After meeting FR1 criteria, mice were moved to FR5 testing, where the reinforcement schedule requires five pokes to earn 1 reward, for three consecutive days.

For PR testing, mice were tested on a progressive ratio schedule of reinforcement where the number of nose pokes required to elicit reinforcement is calculated using the equation Reinforcements = ‖5e^(N*0.2)^‖ – 5, where N is equal to the number of sucrose solution reinforcements already earned plus 1. When mice earned the same number of rewards (+/−10%, or within 1 if less than 10 rewards are earned) for 3 consecutive days, they were considered stabilized at their “breakpoint”. For full methods and instructions, see published online methods at https://zenodo.org/record/1011360 ^28^.

### Statistical analysis

All data were processed using R statistical software ^57^. Welch’s *t*-tests were used to correct for unequal variances, linear mixed effects ANOVAs were run using the ‘nlme’ package, effect size was calculated with the compute.es package ^59^, and power was calculated using the pwr package ^60^. Graphics were produced using ggplot2 ^61^ and Illustrator (Adobe).

## Acknowledgements

This work was supported by funding from the Western Washington University Behavioral Neuroscience Program. The authors would like to thank Dr. Janet Finlay for sharing her protocol for reducing mouse body weight in preparation for the behavior experiments, and Jim Mullen and Jason Bryenton for technical assistance with the design and building of isolation housing.

## Competing financial statement

The authors declare no competing financial interests.

## Author contributions

SM and RMB worked together to devise the experiments, modify the behavioral apparatus, analyze the data, and prepare the manuscript and should be considered co-first authors. SM, HST, WTS, EMV, MH, SRWL, DDS, and SRC collected the data for the initial behavior cohort. HST, WTS, RMB, and SM planned and collected the data with WSH for the replication cohort. JPC and JBC planned and supervised the experiments.

---

**Supplemental Fig 1.**
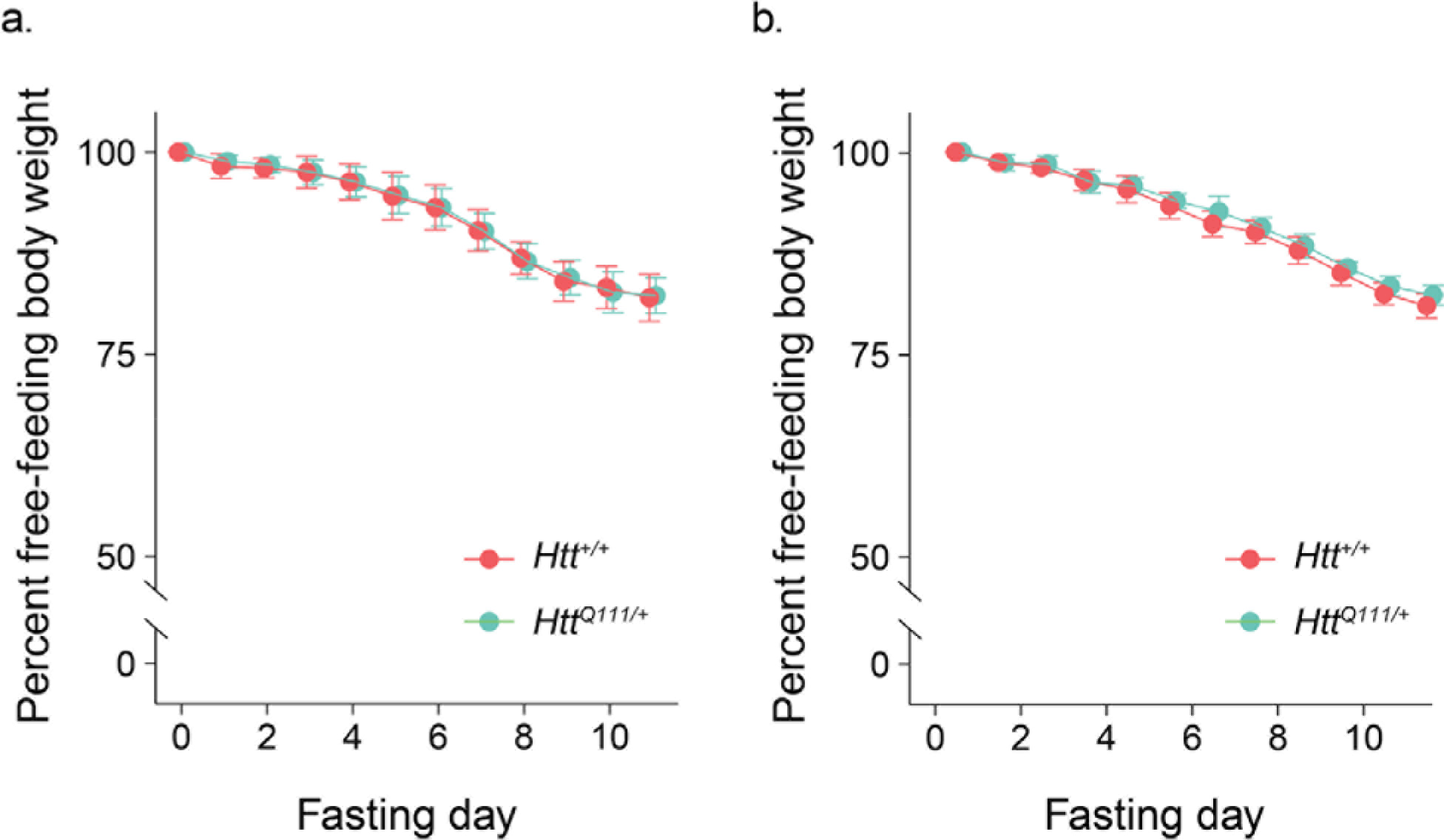
Body weight during food restriction of 10-month-old *Htt*^+/+^ and *Htt^Q111/+^* mice. Body weight was reduced during 11 days of food restriction, but genotype did not change overall body weight, or the rate at which mice lost weight . **a.** Initial cohort: linear mixed effects model, effect of date *F*_(11, 187)_ = 673.6, *p* < 0.0001; effect of genotype *F*_(1, 17)_ < 0.001, *p* = 0.96; genotype x date interaction *F*_(11, 187)_ = 0.5, p = 0.87). **b.** Replication cohort: linear mixed effects model, effect of date *F*_(11, 242)_ = 889.9, *p* < 0.0001, effect of genotype *F*_(1, 22)_ = 2.6, *p* = 0.12, effect of genotype x date interaction *F*_(11, 242)_ = 1.33, *p* = 0.21. Error bars represent S.E.M.

